# Experimental insights on the behaviour and development of Near Eastern Fire salamanders (*Salamandra infraimmaculata*) indicate general strategies to cope with aquatic predators

**DOI:** 10.1101/2025.02.07.637136

**Authors:** Timm Reinhardt, Eliane Küpfer, Ori Segev, Leon Blaustein, E. Tobias Krause, Sebastian Steinfartz

## Abstract

<u>F</u>ishes are key predators of amphibian larvae with often devastating consequences for their abundance and survival. <u>In Israel, t</u>he Near Eastern Fire salamander (*Salamandra infraimmaculata*) reaches the southern limit of the geographic distribution of the whole genus <u>and has to</u> cope with xeric terrestrial habitats- and a scarcity of suitable sites for the deposition of larvae <u>with permanent</u> springs and streams are <u>being</u> rare. One such location is the Tel Dan National Park in the North of Israel, were, surprisingly salamander larvae and fishes coexist successfully in a system of larger and smaller streams. In three experiments we tested different anti-predator behaviour with salamanders from Tel Dan and other locations. We analysed <u>w</u>hether gravid females avoid the presence of fish when depositing their larvae; if the presence of fish impacts larval development and if salamander larvae show active hiding behaviour when confronted with fish cues. While females from Tel Dan showed a bet-hedging strategy in site selection irrespective of fish presence, larval developmental rate was reduced in the presence of fish probably due to a lower feeding activity. Lastly, larvae from different sites displayed hiding behaviour in the presence of fish, irrespective whether they naturally coexist with fish or not. We infer from our results and existing studies that *S. infraimmaculata* has general rather than site-specific adaptations to cope with the presence of fish. We conclude that the general adaptations are essential to sustain this species in a range of extreme habitats at the southern limit of the s distribution.

## Introduction

Among tetrapods, amphibians are classically associated with freshwater habitats as their simple eggs and larvae are barely protected from dehydration and desiccation. Consequently, the reproduction of amphibians is directly or indirectly associated with fresh water and many amphibian larvae spend their time until going on land (i.e. completing metamorphosis) in freshwater habitats. In such habitats, however, interspecific interactions, such as predation and competition, with fishes, the other group if aquatic vertebrates, are important biotic factors limiting the occurrence and distribution of amphibians. Fishes are top-predators of amphibian eggs, larvae, and even adults usually with devastating impacts on amphibian populations and communities (Hecnar and M’Closkey, 1997; Hartel et al., 2007; Segev et al., 2009). In experimental setups, when fish had been removed and excluded from the downstream regions of streams, larvae of *Ambystoma texanum* survived in such regions of streams where they usually did not occur. Conversely, by introducing fish to amphibian breeding ponds, salamanders were extirpated (Petranka,1983). Besides being important predators, fish may also compete with the amphibian larvae for food and spatial resources (Blaustein and Chase, 2007). While fishes are normally strictly dependent on the permanent availability of water, amphibians with their biphasic (aquatic and terrestrial) life cycle often reproduce in ecologically unstable, small, and ephemeral water bodies which are typically unsuitable for permanently aquatic species, thereby reducing interaction with fishes (Werner and McPeek, 1994; Hamer and Parris, 2013). In general, amphibian species that are sharing the aquatic habitat with fishes have developed specific adaptations in their evolutionary history to minimis or avoid predation (Sih et al., 2003).

One such specific response to predation is brood-care behaviour of the mother (e.g. Wells, 2007); in its most basic form, reproducing females actively choose the aquatic habitat where to deposit their offspring to increase their survival. Females might select sites with lowered risk of predation for their offspring. Avoidance behaviour — where amphibians do not use water bodies where fish are present — has been shown in many species including salamanders (e.g., Storfer et al., 1999; Rieger et al., 2004;).

Another general strategy to avoid predation is performed by the offspring themselves; predator avoidance mechanisms of amphibian larvae can be either based on morpho-physiological or behavioural adaptations. Morpho-physiological adaptations of amphibian larvae can include colour changes for better camouflage (Kats and Sih, 1992; Segev, 2009, van Buskirk and Schmidt 2020) or change in body size to decrease attractiveness or susceptibility to predators (Preisser and Orrock, 2012). The general tendency to escape unfavourable conditions, e.g. risk of desiccation but also predation by the reduction of time until metamorphosis is an important mechanism to avoid predation (e.g. Relyea, 2007) but also costly as it usually goes along with a reduced size at metamorphosis as discussed by Semlitsch et al. (1988) and Weitere et al. (2004). Behavioural responses of amphibian larvae to predator presence include also increased “shyness” — which would be reflected in a reduced activity in the open, an increase in refuge usage, and change in diurnal activity (Semlitsch, 1987; Watson et al., 2004, Petrovic et al., 2020). These individual predator responses by amphibians are typically triggered by cues, such as the visual recognition of a predator, or the olfactory perception of kairomones, which are substances exuded by fish when present in an aquatic environment and perceived and used as alarm cues by potential prey species (Schoeppner and Raleyea, 2009; Mathis et al., 2003). Similar alarm responses can be triggered by pheromones, secondary metabolites or body fluids originating from conspecifics (Hews and Blaustein, 1985).

Fire salamanders are members of the clade of true salamanders (Salamandridae) and species of the Genus *Salamandra* are distributed across major parts of Middle and Europe (*Salamandra salamandra*), the island Corse (*S. corsica*), parts of Northern Africa (*S. algira*). In the Near east, *Salamandra infraimmaculat*a; (Iran, Irak, Syria, Lebanon and Israel see Vaissi 2021 for details) occupies the southernmost limit of the range of the genus Salamandra (Steinfartz et al., 2000, Vences et al., 2014, Goedbloed et al., 2017). Not only among urodelans (tailed amphibians) but also among all amphibians, fire salamanders are unique regarding their reproductive biology. The eggs are internally fertilized by the sperm from sometimes several males (see Steinfartz et al. 2006, Caspers et al. 2014). Embryonic and early larval development occurs in the oviduct. Females select suitable aquatic habitats for the larvae development which are still encapsulated when deposited and “hatch” from the membrane upon touch with water. Most fire salamander species are larviparous (Greven 1998) with a few and populations of *S. salamandra* being viviparous, i.e. giving birth to fully developed juveniles (see Alarcon-Rios et al. 2019, Velo-Anton et al. 2012). Fire salamanders (*Salamandra salamandra*) in central Europe normally occur in lower to middle mountainous regions where fish-free permanent small first order streams and springs are abundant. Here, females have several opportunities for deposition sites for their larvae. Accordingly, the co-occurrence of larvae of *S. salamandra* with fish as predators is very rare and often a result of catastrophic drift events that had transported larvae from the small spring region to larger upstream regions (Reinhardt et al., 2017; Schafft et al., 2022). This downstream drift into the fish region is almost always fatal for the larvae. The presence of fish typically excludes the presence of salamander larvae (Thiesmeier and Schuhmacher, 1990).

Fire salamander species occurring in the southern Mediterranean region, such as *S. infraimmaculata* in Isreal are mainly active during the colder and wet season starting in late autumn (November) until early spring (February). For the rest of the year, they face dry and hot climatic conditions, which are unsuitable for daytime activity. Suitable sites for reproduction are scarce, and the availability of water is constrained to the short-wet season of the year. Often larval habitats are stagnant (ephemeral) water bodies, many with short hydroperiods. In Israel, livestock troughs and rock pools, which are filled by ephemeral stream are typical reproduction habitats (Zacharias et al., 2007). The deposition and development of larvae in permanent streams is an exception (Degani, 1996; Goldberg et al., 2009; Blank and Blaustein, 2014). In contrast to populations of S. *salamandra* in Europe, where larvae can occupy fish-free streams or regions of a stream, larvae of *S. infraimmaculata* have no alternative but to co-exist and cope with different fish species in certain permanent streams. The Tel Dan region is a unique ecosystem in Israel with many smaller and major streams building the main sources of the river Jordan. A dense and humid mediterranean broad-leaf forest comprises the terrestrial habitat. Indeed, the Tel Dan population is one of the largest salamander populations known for Israel (Segev et al., 2010). Females have a multitude of opportunities to deposit their larvae in one of the permanent rivulets. In these, however, an important drawback is the presence of predatory fish in the stream system. As documented by their high population size, however, fire salamanders must have developed certain strategies to minimise or avoid encounters with fishes. One observable behavioural adaptation is the strong hiding behaviour of the larvae under stones in the interstitial which is in contrast to the behaviour of their European counterparts which grow up in fish-free streams.

This study aims to investigate in how populations of *S. infraimmaculata* – with a special focus on the Tel Dan population - show traits representing a specific avoidance behavior in response to fish as potential predators on their larval. During our study, we address the following questions i.) Do females of *S. infraimmculata* avoid water bodies with fish presence when depositing their larvae ii.) Do the larvae respond to the threat of fish predation physiologically by accelerating their developmental rate and thus time to metamorphosis iii.) Do larvae also react to the predation threat with increased shyness and hiding behaviour.

Finally, we want to investigate, if these responses are part of a general avoidance strategy of *S. infraimmaculata* populations or specific for populations that naturally coexists with fish as in the case of the Tel Dan population.

## Material and Methods

To address these hypotheses, we designed a series of experimental setups involving adults and larvae of *Salamandra infraimmaculata*. The animals originated from various sites in Israel that differed regarding the natural co-occurrence with fish and salamander larvae. The first experiment (larviposition choice experiment) tested if adult females of *S. infraimmaculata* discriminate between pools with or without fish presence for the deposition of their larvae. In the second experiment, we tested in how far larval development and metamorphic timing was influenced by the water-borne fish cues that mimic the presence of predators. In a third experiment we tested whether hiding behaviour increased in the presence of fish cues.

### Study sites

We included individuals from sites in three regions in Israel, the pond Kaukab in the Galilee Mountains in central Israel (fish presence), the ponds Secher and Ein Al Balad (referred to as Balad in the following) on Mt. Carmel in the Southwest of Israel (fish free) and from streams of the Tel Dan National Park in the Northeast of Israel (fish presence) (Table 1).

**Table 1:**
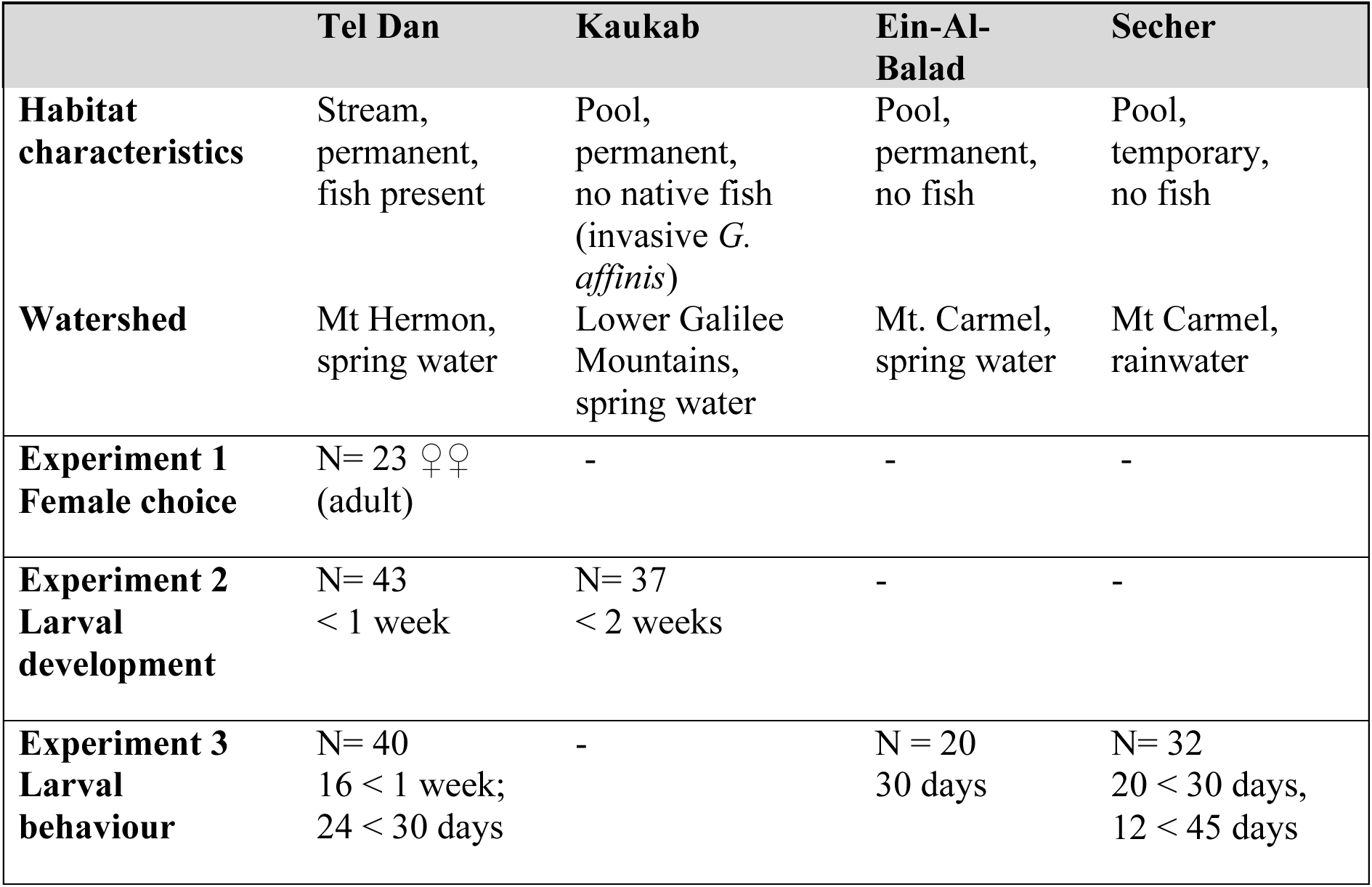
Summary of experimental design parameters and the sites of origin of the 23 female *S. infraimmaculata* used in experiment 1, and overall (N=172) larvae used in the experiments 2 and 3.

All pond sites are temporary pools filled with rainwater or spring water (so called “Wadis”). They experience large temperature fluctuations between 14°C in winter to more than 30°C in the hot season. The Ponds Secher and Balad are rock pools that are rainwater fed and flushed occasionally and naturally free of fish. The pond Kaukab is permanently spring fed and harbors the mosquitofish (*Gambusia affinis*). Prey items for salamander larvae in these pools include mosquito larvae (mostly Culicidae), Cladocerans and Ostracods.

Fire salamanders occur along an elevational gradient in the Dan-Jordan River system, from the floodplains to the headwater region and spring locations in the Golan heights (see Preißler et al., 2020). The Tel Dan Nature reserve is one of the few permanent limnic systems in Israel inhabited by *S. infraimmaculata*. The salamander larvae are found in the springs and rivulets that were formed by permanent karstic springs. The springs are fed by rain and snowmelt from Mt. Hermon and the water has a constant temperature of 14.5 °C throughout the year (Gil’ad and Bonne, 1990). Potential invertebrate prey for salamander larvae found in the Tel Dan streams includes freshwater amphipods (*Gammarus syriacus*), several species of mayflies (cf. *Ecdyonurus spp.*), snails (*Melanopsis sp.*) and the larvae of non-biting midges (Chironomidae). Other invertebrates are potential predators of the salamander larvae, such as the backswimmer *Notonecta sp.,* leeches (Erprobdellidae), and freshwater crabs (*Potamon potamios*). Small cyprinid fish, such as the Levantine Minnow (*Pseudophoxinus kervillei*) and the stone loach (*Oxynoemacheilus insignis*) can be found in deeper parts of the rivulets in syntopy with the salamander larvae. Downstream, the river system is occupied by larger cyprinid fish such as the Levantine Scraper (*Capoeta damascina*) – a native barb species, and the introduced North American rainbow trout (*Oncorhynchus mykiss*).

### Origin of female salamanders and larvae

Potentially pregnant females were collected from Tel Dan and Kaukab and were kept in terraria (40 x 100 cm ground size), equipped with soil, bark, and leaves to provide shelter as well as a water dish for larvipositioning. Larvae from Kaukab were born under these husbandry conditions at the University of Haifa and the females were subsequently released at their capture site. Most of the females from Tel Dan were transferred to the experimental enclosures (see below) for the deposition of larva. In subsequent experiments we used larvae obtained from these females as well as larvae directly collected from Kaukab, Balad and Secher (see table 1).

### Experimental setups

#### Larvipositioning choice experiment – experiment 1

The aim of this experiment was to test whether salamander females avoid depositing larvae into pools with fish presence. This experiment was conducted with 23 potentially pregnant females collected from the Tel Dan population in an outdoor area of a local fish farm in Kibbutz Dan, adjacent to the Tel Dan Nature reserve. The females had a mean length (SVL) of 123.0 mm ranging between 102.7mm and 147.4mm and a mean weight of 56.92g (ranging between 39.30 g and 78.29g) before larvipositioning. Experimental setup as described in Sadeh et al., (2009) and Segev et al., (2011), we used ten identical walled enclosures (3 x 2m); each enclosure contained four plastic pools for females to deposit larvae (hereafter referred to as pools) with the dimensions of 40 x 50 x 20 cm (length x width x height) for larval deposition (Fig. 1 A). The pools were each filled with 40l of spring-water which was kept at a constant level. We added a similar amount of volcanic rock substrate from the Tel Dan region to each pool to facilitate the females in entering for larval deposition, and dried leaves to provide structure and places for shelter. In the centre of each enclosure, hiding places of dead wood and rocks were created (Fig. 1 A). In each enclosure one salamander female was placed; females could choose between two fish free control pools and two pools containing caged juvenile carp (*Cyprinus carpio*). We used living carp due to their availability at the fish farm and hardiness under the experimental conditions. Most fish species in the Tel Dan rivulets occurring in syntopy are also cypriniforms but could not be used here due to conservation and animal welfare concerns. Cyprinids are known to trigger strong responses in a range of aquatic species (e.g. Hahn et al., 2019). The juvenile carp were about 5-8 cm in length, thus similar in size as the fishes cooccurring naturally in Tel Dan.

**Figure 1:**
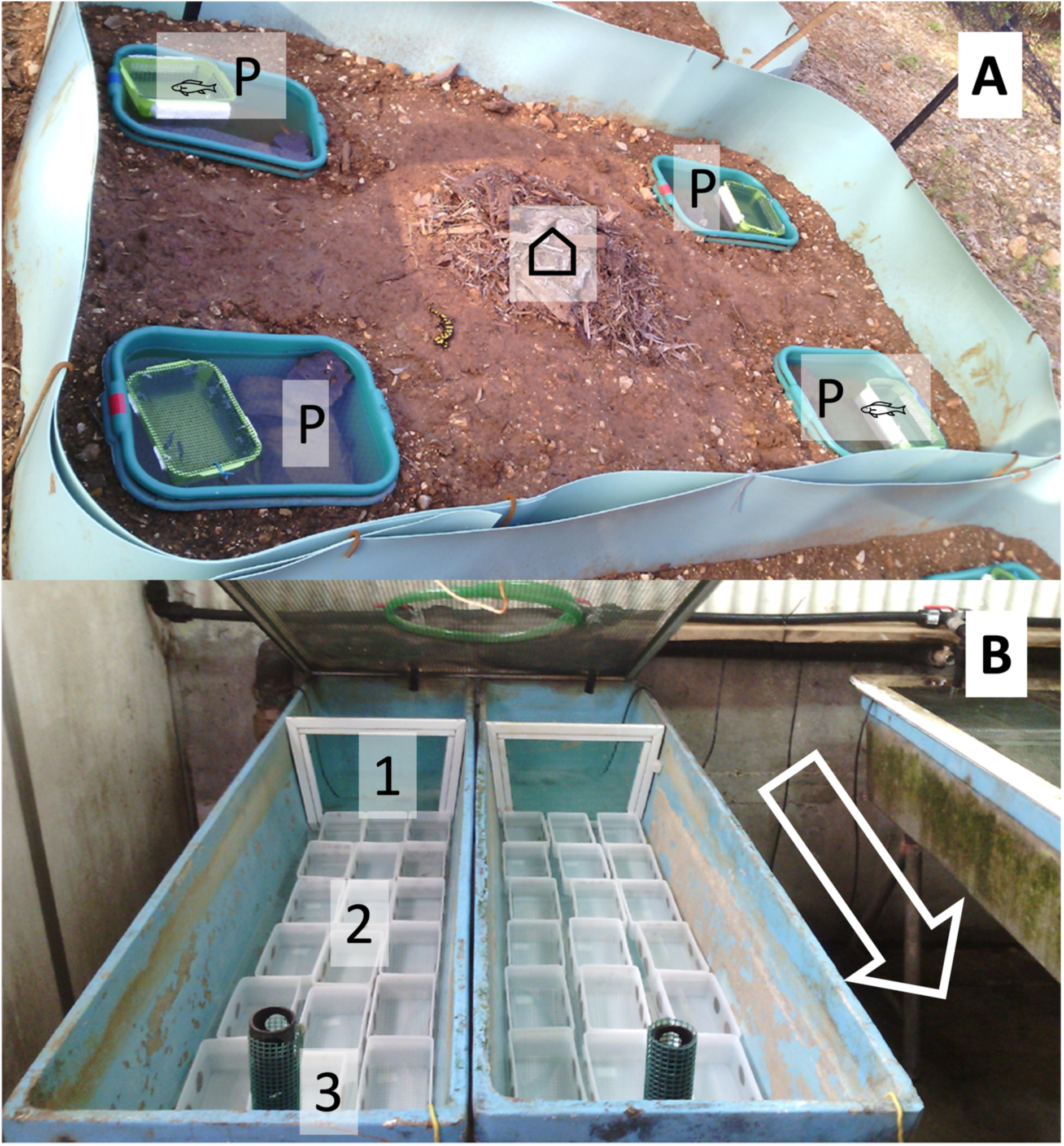
Experimental setups **A)** Larviposition choice experiment (Experiment 1): Each arena (200 x 300 cm) contains (P) four pools (40 x 50 x 20 cm), two stocked with caged young carp (*Cyprinus carpio*) and two empty controls, positions randomised with each trial. In center (⌂) retreat for the *S. infraimmaculata* females. **B)** Setup of larval development experiment (Experiment 2): Two flow channels with (1) holding units for rainbow trout (*Oncorhynchus mykiss*), or no-fish-control at water inflow, (2) separate holding units for 18 salamander larvae and (3) outflow (Arrow indicates direction of water-flow).

In order to present chemical as well as visual predator cues for the depositing females, fish were housed in cages (20 x 25 x 10 cm of plastic mesh, hole size 4 x 4 mm) each containing two fish. Identical but empty cages were added to the control treatment (i.e. the pools without fish). The position of the cages was randomized for each trial.

The fish were removed from the experimental setup during the day — when salamander females were inactive and hiding and no deposition of larvae had to be expected — and housed in separate aquaria where they were fed with frozen chironomid larvae. The salamander females were placed into the enclosures in November 2014 and remained within the enclosure for up to four weeks until they experienced at least one heavy rain event or until they finished deposition of larvae. Salamander females were active during the night and each morning all larvae collected and counted from the artificial pools for subsequent experiments. All female salamanders were released at the site of collection in the Tel Dan reserve after the experiments.

#### Larval development and metamorphic timing experiment – experiment 2

In this experiment we tested how the presence/absence of fish influenced larval development and metamorphic timing in a factorial design. We obtained larvae used for this experiment from gravid females of the Tel Dan population and from females collected near Kaukab, the permanent lentic spring in the Lower Galilee. All larvae were naïve to fish presence. We mimicked lotic stream conditions and constant exposure to fish cues in large flow channels; salamander larvae were placed individually in plastic boxes with mesh windows (10 x 20 cm, 10 cm deep) that allowed fish cues to flow through the boxes but prevented a visual detection of the predator by the larvae (Fig. 1B). As living rainbow trout (*Oncorhynchus mykiss*) were maintained and bred at the fish farm we used this species for the source of emission of predatory fish cues as rainbow trout are known to be predate on salamanders (Hartel et al. 1997). Boxes containing one salamander larva each were arranged in four distinct flow channels, 189 cm long and 34 cm wide with a water level of 9 cm. Each flow channel contained 20 boxes with larvae and a separated compartment housing a trout. The water flow through the channel by entering first the compartment of the fish and then passed the boxes of the salamander larvae before leaving the channel by a sink at the end of the flow channel. Of the 20 boxes in each channel, 10 contained larvae from Tel Dan and Kaukab, respectively. The channels were fed with water directly from underground springs with a flow velocity exchanging the whole water volume once per hour. Fish were present in two flow channels, whereas the other two channels were used as control experimental settings without fish; in total we raised 80 larvae under these conditions until metamorphosis (Table 1). Larvae from both localities (Tel Dan and Kaukab) were distributed in equally to treatment and control channels and randomly placed within each channel. Both the fish and salamander larvae were fed ad libitum with commercially available live *Chironomus spp*.

The wet weight (g) and snout vent length (mm) of each salamander larva was measured every two weeks. Salamander larvae were removed from the experiments as soon as they showed signs of metamorphosis, indicated by the reduction of gills, change of head morphology and coloration pattern (see Sanchez et al. 2018 for a description of larval stages).

The daily growth rate (DGR) was calculated between the age of 14 and 42 days, as follows:

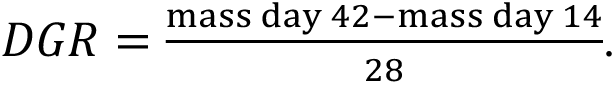

Within this age range, the larvae were still in a phase of approximately continuous linear growth (see also Reinhardt et al. 2013) and did not yet show signs of metamorphosis. After further 30 days, surviving larvae and juveniles (78%) were released in the location where their mothers were collected.

#### Larval hiding behaviour experiment – experiment 3

In this experimental setup we analysed the behaviour of salamander larvae in the presence of fish cues. We tested in total 92 larvae from Tel Dan and larvae of approximately the same size collected at the sites Secher and Balad (see Table 1). The larvae obtained from Tel Dan, Balad and Secher were younger than 30 days but showed differences in size and age. In addition, we tested newly deposited larvae from Tel Dan and larvae from Secher at an age of more than 45 days (Table 1). None of the larvae had been used in fish treatments in any of the previous experiments. The behavioural experiments were conducted in a room with an artificial 12:12 hour day-night cycle. The larvae were set up individually in plastic containers (same type as in experiment 2) containing 5 l of water and a water level of 4 cm. Each setup contained two grey plastic tubes (8cm long, 3cm diameter) as hideouts for the larvae. All experimental containers were cleaned with alcohol and the water was exchanged after each experimental run. For the fish treatment, the water was taken from a 100 l aquarium containing ten rainbow trout.

For each run trial, a single larva was introduced in the centre of the container. Four experiments were conducted simultaneously with two individuals under fish cue treatment and two as control without fish cues. The behaviour was recorded on a video tape for 25 minutes. The first and last five minutes of the recording were excluded and the remaining 15-minute sequence was analysed. We measured if the larvae were hiding in the observational period or not as a binomial response variable (0= not hiding in the observation period, 1 = hiding in the observation period). All larvae survived the experiment and were released in their respective original habitats.

### Statistical analysis

All statistical analyses were performed using R 3.2.3 (R Core Team, 2015). In the larvipositioning choice experiment (experiment 1) a one sample t-test was used to analyse whether deposition of larvae by individual females differed from a 50:50 (fish-presence *versus* fish-free pools) distribution to be expected if females do not discriminate for the presence of fish. Furthermore, we used a Linear Model (LM) with the following explanatory factors i) female condition (as weight/size ratio); ii) total number of larvae deposited; iii) the number of deposition events per female. Residuals of the data were controlled for normal distribution using a Shapiro-Wilk normality test.

In the larval development experiment (experiment 2), we used linear mixed effect models (LME) to compare the daily growth rate (DGR) between larvae from the different sites and treatments and their interaction using and included the identity of the mother as a random factor. The model of the weight at day 14 showed a deviation from normal distributed residuals, due to two outliers which were removed to achieve a normal distribution of the residuals. As a result, the significant effects were not driven by residual outliers (as described Caspers et al. (2014b)). The age at metamorphosis was analysed for all larvae that metamorphosed by the end of the experiment. Treatment, site, and their respective interaction weight at metamorphosis were included as factors in another LME with maternal ID as random factor.

A Binomial Mixed-Effects Model (glmer of the lme4 package; Bates et al., 2015) was fitted for the analysis of the hiding-behaviour (experiment 3), i.e. on whether (=1) or not hiding (=0) was shown by individuals; this conservative transformation was made as distribution of times in the test were not suited, even after transformation, for parametric analyses. The treatment, site of origin and their interaction were included as fixed factors as well as larval age; the maternal ID was included as a random factor.

### Ethical Note

Collection of animals and experimental setup was conducted under the permission of the Israeli Nature and Park Authorities (INPA) under permit no. 2014/40672. All animals were released unharmed in their original habitats following the experiments.

## Results

### Choice of larvipositioning sites – experiment 1

Altogether, sixteen (69.5 %) of the 23 adult females deposited a total of 580 larvae during 73 events during of the study period. The remaining females either deposited larvae already in the acclimatisation phase under laboratory conditions or did not deposit larvae at all.

In 35 larviposition events, 276 larvae were deposited in pools with fish cues, and in 38 events, 304 larvae were deposited in pools free of fish cues (i.e. control pools). The average proportion of larvae deposited in pools with fish cues *versus* control pools did not differ significantly from chance level, i.e. a 50% probability to each treatment (one sample t-test, df=15, t=-0.39, p=0.70; Fig. 2). The overall ratio of deposited larvae in the control pools for each female was not affected by the number of deposition events, nor by the total number of larvae deposited, nor by female body condition (LM: factor number of events (deposition days), F_1,10_=0.76, p=0.40; factor absolute number of larvae F_1,10_=0.40, p=0.54; factor female condition F_1,10_=0.04, p=0.85). None of the observed parameters differed significantly between the fish cue treatment and the controls. Instead, larvae were spread over several pools and deposited during multiple nights without a clear pattern.

**Figure 2:**
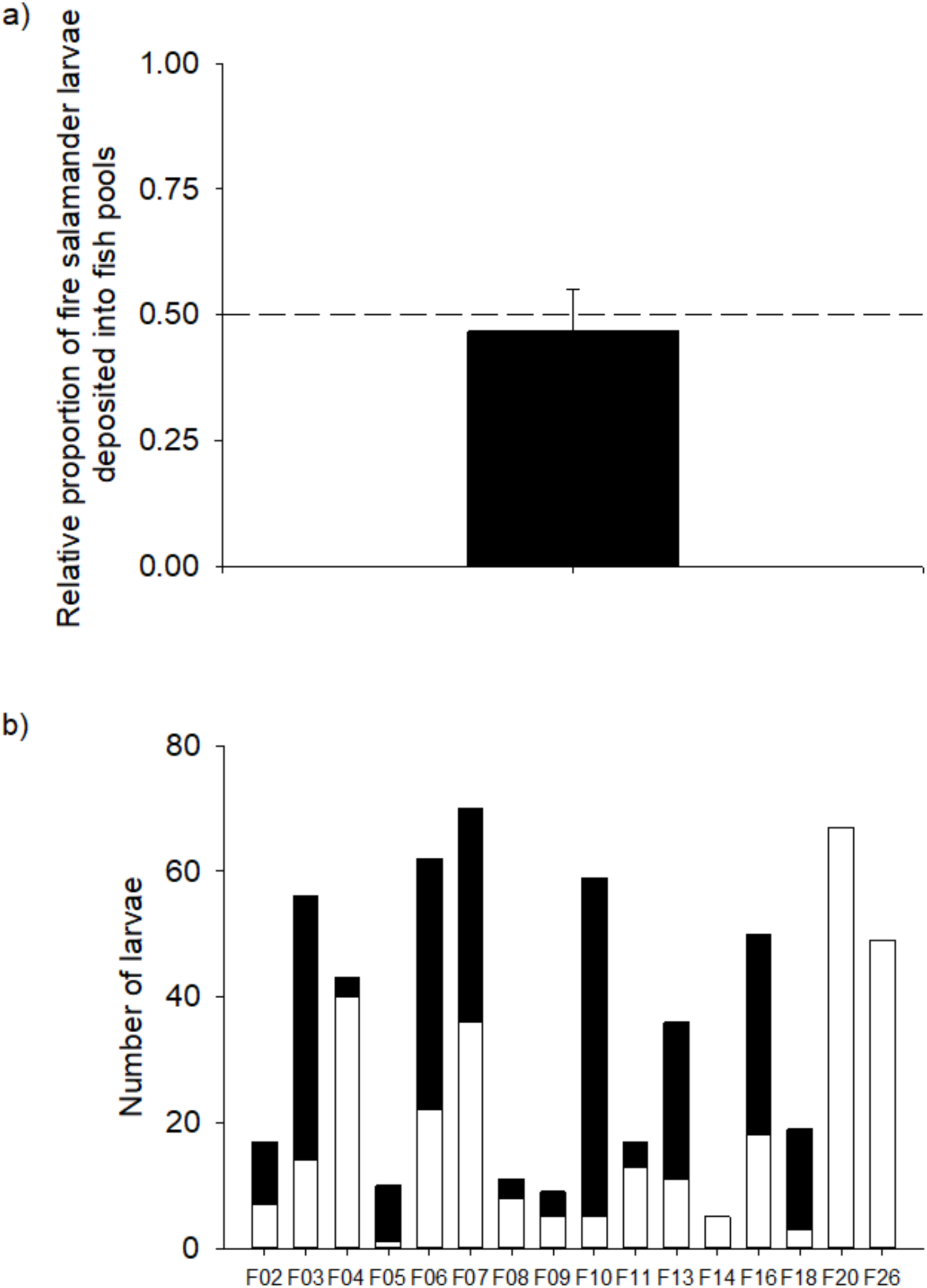
**a)** Relative proportion of salamander larvae deposited into fish-free control pools. Mean ± SE. dashed line indicates the probability level of 0.5. The difference between larvae deposition in fish and control pools was not significant. **b)** Absolute number of larvae deposited in fish-present (black stacks in bars) and fish-free (white stacks in bars) pools of clutches (N=16) of females from Tel Dan population.

### Impact on fish cues on larval development and metamorphic timing – experiment 2

While the mean weight of the larvae after 14 days did not differ significantly among treatments or sites (LME_weight d14_: factor treatment, F_1,64_=3.06, p=0.09, factor sites F_1,6_=0.28, p=0.62; factor treatment*site F_1,64_=0.19, p=0.67), at the age of 42 days, the effect of fish cue presence on larval weight was significant. Individuals from the control (no fish) treatment were on average heavier than larvae from the fish-treatment (LME_weight d42_: factor treatment F_1,66_=8.23, p=0.006; factor site F_1,6_=0.84, p=0.40; factor treatment*site F_1,66_=2.11, p=0.15). Also, the daily growth rate (DGR) between day 14 and 42 was significantly affected by an interaction between treatment and site. The larvae from Tel Dan showed the lowest DGR in the fish-treatment, while DGR was similar in all other treatments (LME _growth rate_: factor treatment, F_1,64_=4.51, p=0.038, factor sites F_1,6_=1.04, p=0.35; factor treatment * site, F_1,64_=4.57, p=0.036, Fig. 3).

**Figure 3:**
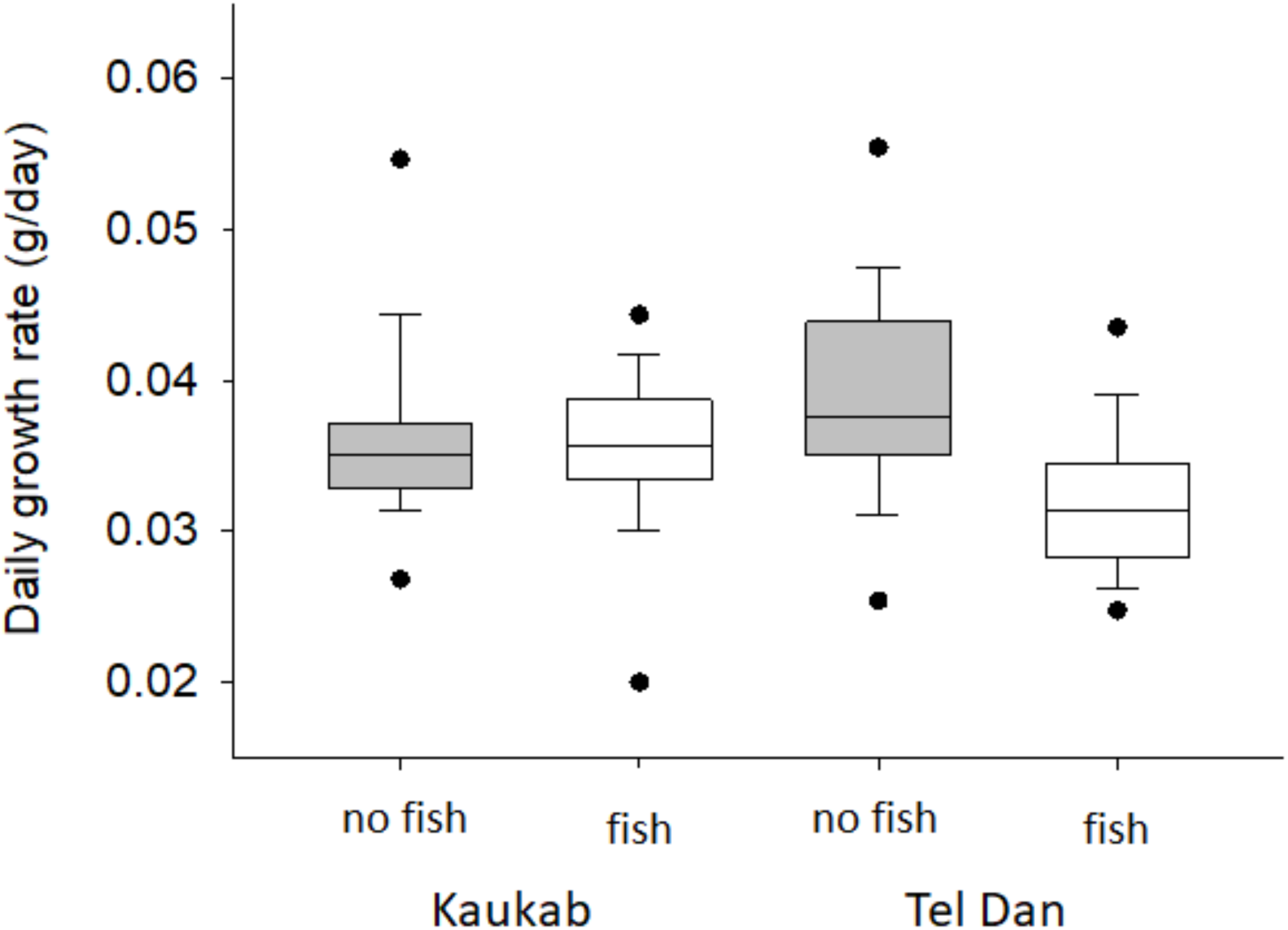
Daily growth of salamander larvae from Kaukab and Tel Dan populations with and without detectable fish cues. The lines in the boxes indicate the median, while the box shows the 25% and 75% percentiles. Whiskers indicate the data range, while the lower and upper dots show the 5 and 95 percentiles.

The age at metamorphosis was not significantly affected by either treatment or site, however older larvae also had a higher weight (LME_age metamorphosis_, factor treatment: F_1,54_=0.01, p=0.91, factor site F_1,3_=0.31, p=0.61; factor treatment*site F_1,54_=0.02, p=0.88, factor weight at metamorphosis F_1,54_=16.01, p=0.0002, Tab. 2). The difference between the weight at metamorphosis between sites and treatments however was not significant (Tab. 2)

**Table 2:** Weight at metamorphosis and time to metamorphosis for *S. infraimmaculata* larvae of the two populations used in experiment 2 in dependence of presence or absence of fish cues. Only the time to metamorphosis was statistically different for the Tel Dan population under fish cue exposure.

| Site | Treatment | N | Weight at metamorphosis<br>(g, mean± SD) | Time until metamorphosis<br>(d, mean± SE) |
| --- | --- | --- | --- | --- |
| **Tel Dan** | Fish | 17 | 1.98 ± 0.33 | 69 ± 2.35 |
| | No fish | 18 | 2.02 ± 0.31 | 60 ± 1.34 |
| **Kaukab** | Fish | 13 | 2.20 ± 0.28 | 68 ± 0.62 |
| | No fish | 13 | 2.38 ± 0.28 | 68 ± 0.80 |

### Larval hiding behaviour in response to fish cues – experiment 3

In total, the hiding behaviour of 92 larvae from three sites was analysed. Overall, we observed a general and strong effect of fish cues originating from trouts on the behaviour of larvae irrespective of site of origin of larvae. When exposed to fish cues 71 % of all salamander larvae went into hiding during the observation, while only 37 % of larvae did so in the control treatment (GLMER_hiding_, df= 84, factor treatment: F_1_ =8.85, p=0.03, factor site F_2_=0.40, p_TD_=0.85, p_Secher_=0.75; factor age F_1_=3.94, p=0.05, factor treatment*site F_2_=0.37, p_trt:siteSecher_=0.40, p_trt:siteTD_=0.60; Fig. 4). On average, larvae from the three distinct sites spent more time hidden when fish cues were present while the tendency of hiding decreased for older larvae compared to younger ones.

**Figure 4:**
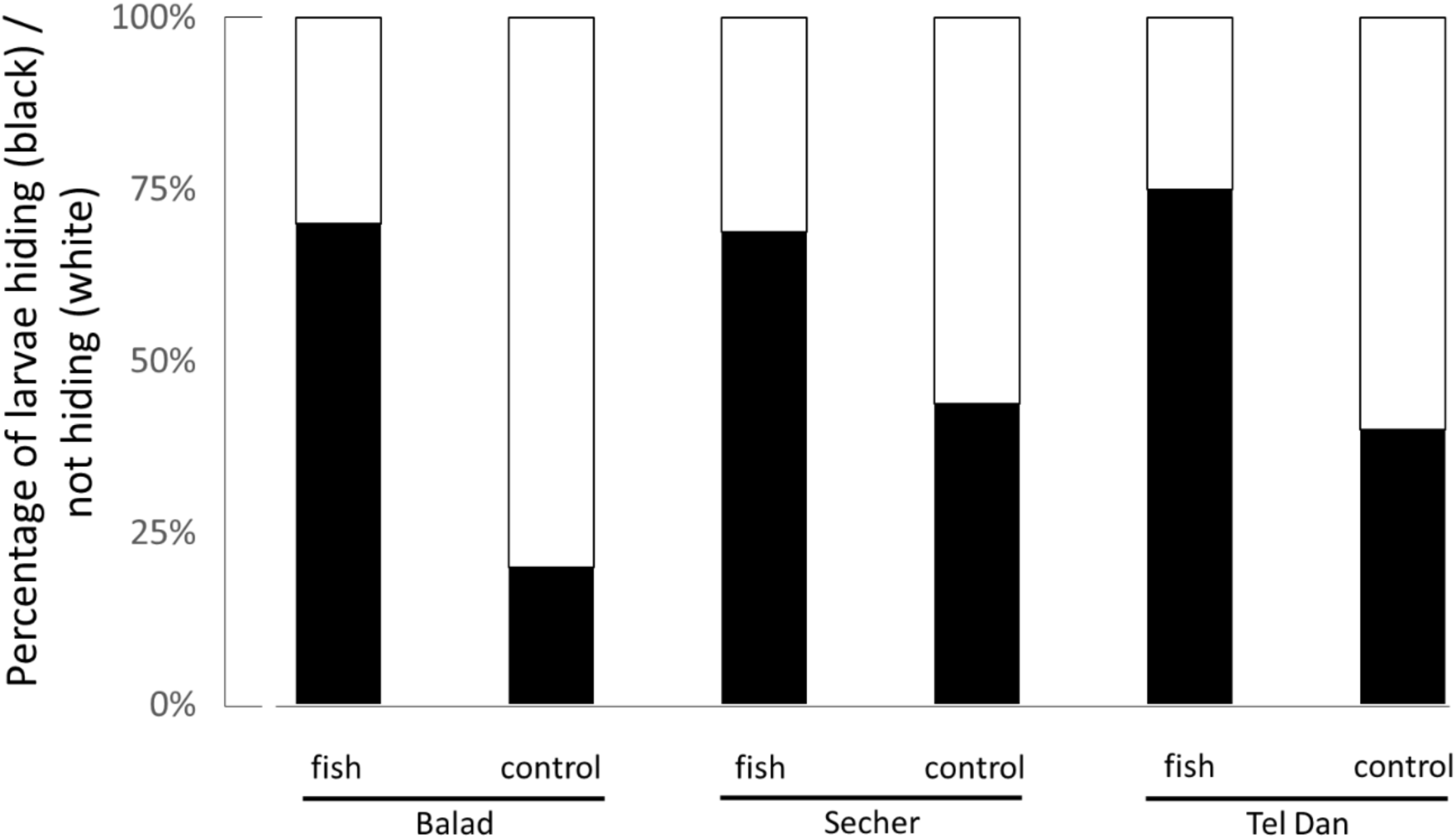
Percentage of larvae that showed hiding behaviour (black stack proportion), respectively no hiding behaviour (white stack proportion) in fish cue treatment vs. without fish cue (control) for larvae originating from three different sites Balad, Secher and Tel Dan. All larvae shown were younger than 30 days.

## Discussion

Our study design was aimed to investigate possible behavioural mechanisms in *S. infraimmaculata* to avoid the co-occurrence of salamander larvae with predatory fish on the maternal and larval level. In the first experiment, we tested whether pregnant females avoid the deposition of their larvae in pools where fish (i.e. fish cues) were present; in the second experiment we analysed whether larval development and/or metamorphic timing were influenced by the presence of fish and finally (experiment 3) we tested larvae for active hiding behaviour in the presence of fish. If having the choice, females do not prefer fish-free aquatic habitats for the deposition of their larvae, whereas specific effects in larval development as well as hiding behaviour can be observed in the presence of fish. Interestingly, these effects are independent whether larvae used in the experiments do naturally co-occur with fish. Therefore, mechanisms to avoid fish at least at the larval stage seem to be a basic underlying behavioural trait in *S. infraimmaculata*.

### Lack of avoidance behaviour towards sites with potential fish predators

Experiment 1 could not support the idea that females avoid waterbodies with fish presence for the deposition of their larvae. The reasons for this are somewhat difficult to assess and further experiments would be necessary. It could be possible that the mothers’ reactions to fish as potential predators are species-specific. In our experimental setup we used common carp which is not native to the Tel Dan reserve where females used in the experiments originated. Species which are not naturally predators might not generate a behaviour of avoidance; a fact that might apply for the fishes living in syntopy with salamander larvae in the rivulets of the Dan River. Or on the opposite, the presence of carp did trigger avoidance behaviour but since the provided larvipositioning sites had no other distinguishable features the females would eventually utilize the pools by chance. In other experimental contexts, *S. infraimmaculata* females distinguished between larvipositioning sites based on structural complexity and the presence of conspecific larvae, which are potentially cannibalistic (Sadeh et al., 2009). In our experimental setup, however, all pools were identical and randomised thus structural complexity was not accounted for.

In fact, all but two females spread larvae of one clutch over several pools or through several subsequent nights. Such behaviour can be best characterized as a bet-hedging strategy, which is well documented for *S. salamandra* in pond environments in Europe (e.g. Caspers et al., 2014a). Bet-hedging strategies in female *S. infraimmaculata* reduces larval mortality which is important as an adaptation to arid and unstable aquatic environments (Segev et al. 2011). It is equally effective at reducing the risk of losing the whole clutch to predation (Lips 2001, Thumm and Mahony, 2002). Bet-hedging can be seen as a general and overall successful strategy in *S. infraimmaculata* (e.g. Sadeh et al., 2009).

Another potential explanation for the lack of avoidance behaviour of the females could be the fact that in almost all smaller and larger streams in the Tel Dan habitat fish are present. Thus an avoidance behaviour would simply make no sense as females cannot choose sites without fish cues. As larvae are fully developed when deposited they are able to immediately escape and the streams offer a magnitude of hiding places for the larvae, especially in the riverbed, a specific female avoidance behaviour to fish cues might not be under strong selection.

### Salamander larval response to fish cues

In our two experiments involving larvae of *S. infraimmaculata* (experiment 2 and 3) we found responses of larvae to the presence of fish cues. We observed that the larvae responded to the threat of predation physiologically as well as behaviourally; we initially hypothesized that larvae would react by accelerating their larval development to minimize the exposure to predation. However, we found that larvae from both tested populations (Tel Dan and Kaukab) showed a reduction in the mean growth rate in response to fish cues rather than an increase. This decrease was significant only in larvae from Tel Dan (see Figure xx) and larvae gained less weight and needed more time until metamorphosis; yet the mean weight at metamorphosis did not differ between treatments or sites. Typically, developmental acceleration and metamorphosis at a lower weight/ smaller size is discussed as a trade-off between escaping the risky aquatic habitat and a resulting lower fitness after metamorphosis (Semlitsch et al., 1988; Székely et al., 2020). Disadvantages of a smaller size and/or lower mass at metamorphosis in amphibians include increased susceptibility to terrestrial predators, reduced movement distance and different ecophysiology, e.g. the ability to retain water (Thorson, 1955, Preisser and Orrock, 2012). Especially, the latter two aspects might be relevant taking into consideration the dry and xeric habitats of fire salamanders in the Near East.

For *S. salamandra*, the stressors ‘reduced water level’ and ‘food shortage’ triggered metamorphosis, both in laboratory experiments and under natural conditions (Weitere et al., 2004; Reinhardt et al., 2013). Under the absence of metamorphic triggers urodele larvae remain in the aquatic larval stage until they reached an ideal body condition for a life after metamorphosis (Werner 1986). We show here that for *S. infraimmaculata*, fish cues alone are not a sufficiently intense trigger to induce metamorphosis, even when the minimal size threshold has already been reached and or exceeded. Indeed, the habitat features of the Tel Dan streams – which are stable in terms of water level and food availability – support the idea that accelerating metamorphosis might not be necessary for the larvae, especially as larvae can hide in the stream ground preventing predation. In comparison to the permanent but more fluctuating pond Kaukab, Tel Dan larvae can complete their larval development without the need to react strongly to metamorphosis triggers; a trade-off between body size and developmental speed might even be maladaptive. *S. infraimmaculata* is the largest species in the genus Salamandra, with adults growing to more than 30 cm in total length and metamorphlings weighing between 2 - 4 g (Goldberg et al. 2009) For comparison: European *S. salamandra* rarely exceed an adult size of 20 cm and more than 1.7 g at metamorphosis (Reinhardt et al. 2013). The large size is likely an adaptation to the dry terrestrial environment. In this context, a larger size at metamorphosis could be more important than leaving the larval habitat too early due to the presence of potential predators which can be coped with (e.g. by effective hiding behaviour).

### Larval behaviour could explain growth reduction but not differences in growth patterns

Observed reduced growth rates despite ample access to food could be a direct result of larval behaviour in response to the presented fish cues. In experiment 3 almost all larvae reacted to the predation threat with increased hiding, i.e. as an indicator for shyness as predicted. Larvae sought out the shelter at a higher rate when fish cues were present compared to the control treatments without fish cues.

Yet, increasing the time spend in the shelter logically comes at the cost of time for foraging and feeding (Krause et al., 2011, Krause and Liesenjohann 2012). *Ambystoma* larvae reacted to fish presence by reducing their activity in the open water, which ultimately reduced the encounter rates between larvae and their food, effectively decreasing their food intake and ultimately growth (Storfer and Sih, 1998). We assume a similar effect for larvae of *S. infraimmaculata* raised in experiment 2. Additionally, stress caused by the presence of predators might increase the metabolic costs (Barzaghi et al., 2020, Schulte et al. 2024).

Experiments performed with larvae of S. *salamandra* — exposed to cues of predatory alpine newts (*Ichtyosaura alpestris*) — showed similar reactions to such predators. The reaction was population specific and salamander larvae originating from sites with newts in syntopy spend more time hiding than larvae from newt-free sites (Hahn et al., 2022). This is in line with findings from other amphibians which demonstrated that individuals from high-risk environments, that is circumstances in which the probability of dying though predation is increased, react stronger to predator cues (Dempsey et al., 2022). On the contrary, we could not find behavioural responses to be specific to those populations that naturally coexist with fishes. Thus, a difference in hiding behaviour is unlikely to explain the difference in growth observed in the previous experiment. Larvae from all tested sites reacted at similar rates to fish cues. Our findings might reflect a lack of specificity concerning predator cues (Dempsey et al., 2022). The ponds, though naturally free of fish, are certainly not in general predator free (e.g. newts or dragonfly larvae) and reacting to a predator cue would always be adaptive. Moreover, flexible responses to predators in terms of growth and behaviour might be meaningful for inhabitants of seasonal ponds as predators like newts might invade the ponds suddenly while at the same time hiding spots are limited.

## Conclusions

Ecological factors such as predation, hydroperiod, habitat structure and food availability shape a complex system of stressors affecting the Near East fire salamander, *Salamandra infraimmaculata* (Blank et al. 2013). Our study focusing on possible responses to predators suggests a relatively high flexibility of *S. infraimmaculata* to many of these factors based on few traits. The generalist nature of these traits such as a bet-hatching strategy in reproduction, a large body mass at metamorphosis and a generally strong behavioural reaction to cues indicating a risk of predation likely allows *S. infraimmaculata* to use a broad range of (aquatic) habitats across their range.

We assume that there might be little pressure to evolve population specific adaptations in response to predators, as the generalist adaptations are already sufficient. Instead, we need to see the adaptations in a wider habitat context. The dry-xeric conditions and limited availability of breeding sites are likely the more important selective force at the southern distribution limit for the entire genus Salamandra.

## Acknowledgements

We would like to thank Hava Goldstein, Rabea Dabous, and the rangers of the Tel Dan Nature Reserve for their friendly support with the collection of the salamander females. We would also like to thank Avshalom Hurvitz, Arlo Hinckley, Christian Wartenberg, and Daniel Goedbloed for their help and support. Collection of animals and experimental setup was conducted under the Israeli Nature and Park Authorities (INPA) permit no. 2014/40672. This study was funded by the German-Israel Project-cooperation (DIP, DFG reference number BL 1271/1-1, STE 1130/8-1).

